# Patch-Clamp Recording of K^+^ Current from Lacrimal Gland Duct Cells

**DOI:** 10.1101/670653

**Authors:** Loren D. Haarsma, John L. Ubels

**Affiliations:** Departments of Physics and Astronomy, Calvin College, Grand Rapids, MI USA; Departments of Biology, Calvin College, Grand Rapids, MI USA

## Abstract

Electrophysiologic studies have characterized ion channels in lacrimal gland acinar cells, but due to relative paucity and inaccessibility, such studies on lacrimal gland duct cells are challenging. The duct cells are believed to secrete the high level of K^+^ that is present in tears. The goal of this project was to develop a method for isolation of viable, single duct cells, demonstrate their utility for patch-clamp recording and characterize the K^+^ channels expressed by duct cells. Exorbital lacrimal gland slices from Sprague-Dawley rats were incubated with collagenase. Under a microscope, the ducts were microdissected from the glands and then incubated with elastase and collagenase. Dispersed duct cells were plated on a cover slip coated with BD Cell-Tak. Duct cells were distinguished from acinar cells by their smaller size and lack of granularity. Whole-cell K^+^ currents were recorded from duct cells using the perforated-patch technique and pipettes with resistances of less than 10 MΩ. EGTA and CaCl_2_ in the pipette solution were adjusted to give 1 uM free Ca^++^. When held at −80 mV, duct cells showed K^+^ currents that activated at command voltages near 0 mV and reached amplitudes near 1 nA at +100 mV. Currents reached peak amplitude less than 20 ms after depolarization and did not inactivate. These currents were inactivated by holding the cells at 0 mV. Currents were blocked reversibly by TEA in the presence of Ca^2+^, but were not blocked by TEA in the absence of Ca^2+^. Currents were not affected by clotrimazole (10 uM) or Ba^2+^ (5 mM). A method has been established for isolation and dissociation lacrimal gland duct cells for electrophysiologic studies. These cells express a voltage-activated K^+^ channel that is dependent on the presence of intracellular Ca^2+^ and may correspond to the IK_Ca_1 channel expressed on the apical membranes of lacrimal gland ducts.

## Introduction

Lacrimal glands secrete the water and electrolytes of the tear fluid which has an elevated concentration of K^+^ (20-25 mM) compared with other extracellular fluids (4-5 mM),^1,2^ and we have reported that elevated extracellular [K^+^] can protect the corneal epithelial cells from UVB induced apoptosis.^3–5^ Lacrimal gland duct cells are believed to secrete the relatively high [K^+^] in tears.^6^ Channels and transporters for K^+^ and Cl^−^ have been identified in lacrimal gland ducts by immunostaining, and a model for K^+^ secretion has been proposed based on the arrangement of these proteins.^7^ The secretory properties of intact ducts have been described, and the data are consistent with the proposed model.^8–10^

Electrophysiologic studies have characterized several K^+^ and Cl- channels in lacrimal acinar cells,^11–13^ however, due to their relative paucity and inaccessibility, the function of ion channels of duct cells has not been studied extensively. The goal of this project was to develop a method for isolation of viable, single duct cells and demonstrate their utility for patch-clamp recording by characterizing the properties of K^+^ channels in these cells. Electrophysiologic identification of channels in the duct cell membrane will confirm immunohistochemical identification of the channels and support proposed physiologic models of duct function.

## Methods

To isolate duct cells from rat lacrimal glands, male Sprague-Dawley rats between 6 and 8 weeks old were euthanized with sodium pentabarbitol. The exorbital lacrimal glands were removed, dipped in betadine and then rinsed in sterile Hanks BSS. All work with animals conformed to the ARVO Statement for the Use of Animals in Ophthalmic and Vision Research.

Our method for isolating duct cells was developed based on a method used by Tóth-Molnár et al,^8^ who dissected lacrimal gland ducts but did not disperse the cells. The glands were injected with a solution of DMEM with bovine serum albumin (BSA, 1 mg/ml) and collagenase (100 U/ml), and cut lengthwise into four or five pieces. This tissue was then incubated in a shaking water bath with the same enzyme solution for 45 minutes at 37 °C.

Following the first incubation, the tissue was rinsed with a solution of DMEM with BSA (6%) and soybean trypsin inhibitor (STI, 0.1 mg/ml). Under a dissecting microscope, the pieces of lacrimal glands were dissected using microforceps, a 28 ga needle and 3 mm Vaness scissors (Fig. 1a). During the dissection as much of the connective tissue surrounding the ducts as possible was removed. As the ducts were dissected they were stored in ice cold DMEM with BSA (6%) and STI (0.1 mg/ml).

**Fig. 1.**
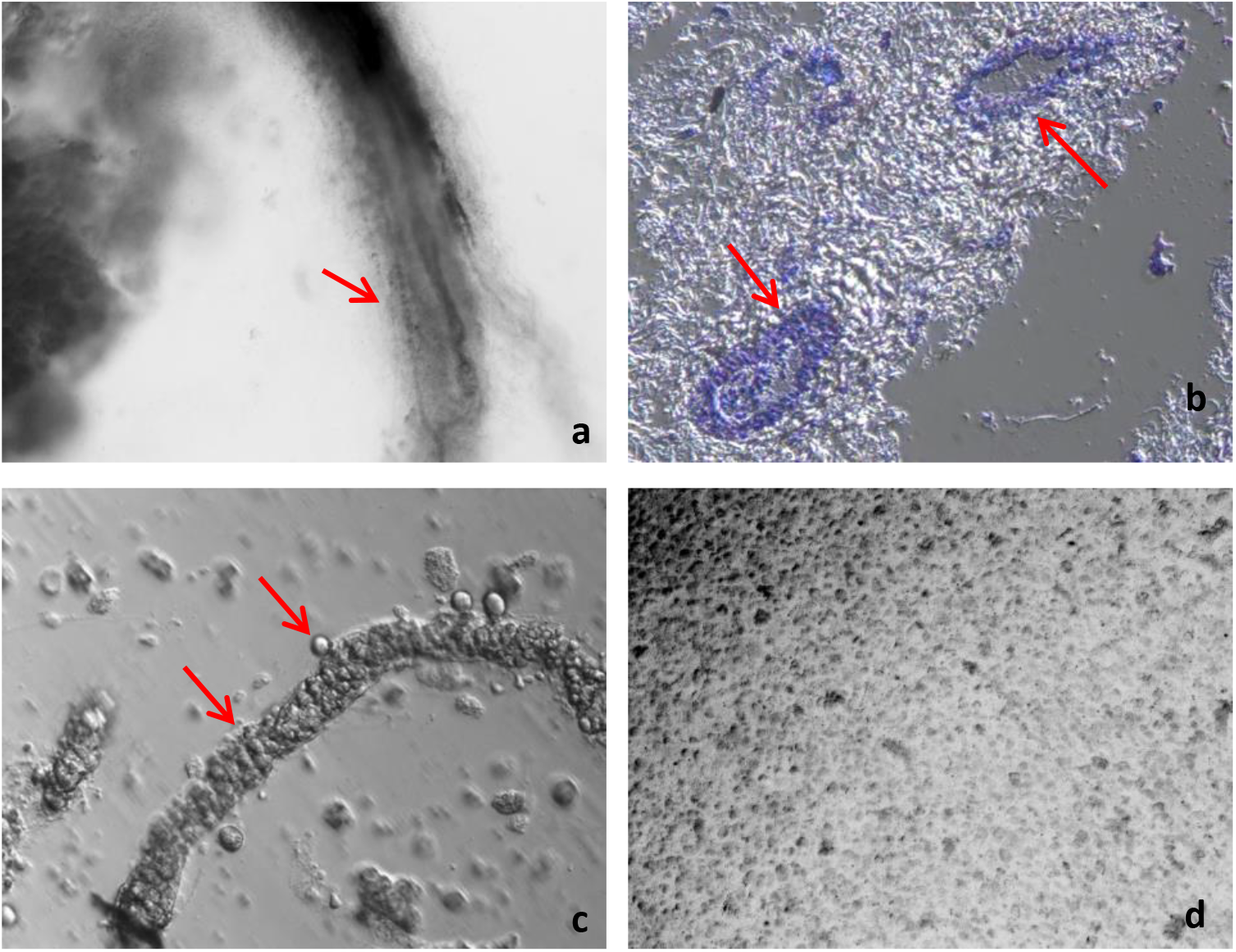
a) Isolated duct after dissection from lacrimal gland tissue. Note the connective tissue still attached to the duct (arrow). b) Frozen sections of dissected lacrimal gland ducts were stained with toluidine blue O to show that ducts were present in the connective tissue (arrows). c) After a 50 minute digestion, some ducts are still intact and some cells have dispersed. Note the lack of connective tissue attached to the duct. d) Duct and acinar cells that have been plated onto a 13 mm plastic cover slip.

After the ducts were removed from the lacrimal gland tissue they were incubated at 37°C in a solution of DMEM with collagenase (300 U/ml) and elastase (150 U/ml) for 50 minutes to remove connective tissue (Fig. 1c). The ducts were then triturated using a Pasteur pipette and incubated for an additional 30 minutes at 37 °C to disperse the cells into a single cell suspension (Fig. 1d).

The collagenase and elastase solution was removed by centrifugation and replaced with 37°C DMEM. The solution containing the remaining duct fragments and cells was then plated onto a 13 mm cover slip coated with BD Cell-Tak. Some acinar cells that adhered to duct fragments were present in the preparations, allowing comparison of ion currents in the two cell types. The cells were incubated on these cover slips for 90 minutes.

For recording, external bath solution (in mM, 160 Na^+^ aspartate, 4.5 KCl, 2 CaCl_2_, 1 MgCl_2_, 5 HEPES Na^+^ salt, pH 7.4) was used to gently wash cells off the cover slip onto a 35 mm recording dish. Although the cover slip was coated with BD Cell-Tak, the cells did not adhere strongly. When washed onto the recording dish after exposure to Cell-Tak the cells adhered readily (Fig. 2).

**Fig. 2.**
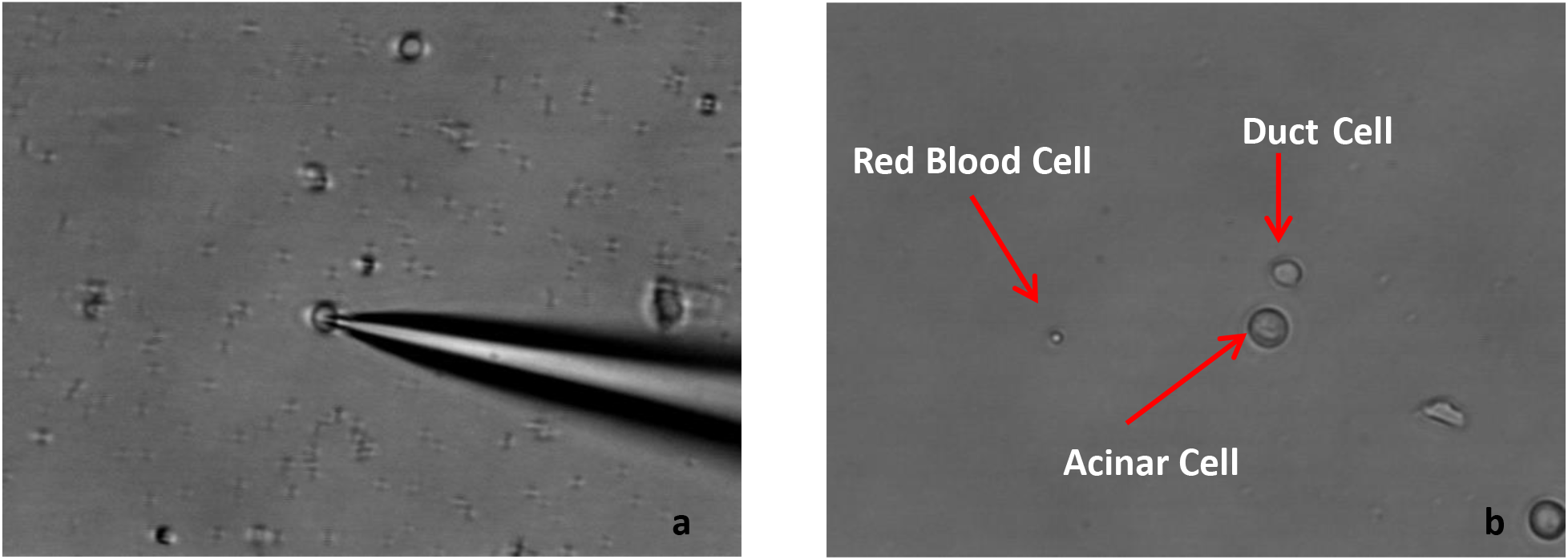
a) Lacrimal gland duct cell with recording pipette attached. b) Acinar, duct, and red blood cells for size comparison. Note granularity of the acinar cell compared with the duct cell. Original magnification 200X

The recording pipettes, which had a tip resistance of 4-7 MΩ to allow attachment to the small duct cells, were filled with internal solution (in mM, 145 K^+^ aspartate, 2 MgCl_2_, 10 HEPES K^+^ salt, 10 K_2_ EGTA, 9 CaCl_2_, pH 7.2) and amphotericin B (250 µg/ml) (Fig. 2). Because Ubels et al. noted Ca^2+^-activated K^+^ channel (IK_Ca1_) staining on duct cell apical membranes,^7^ we recorded using pipette solution both with and without free Ca^2+^. After the access resistance dropped to 60 MΩ or less, depolarizing voltages (−80 mV to 100 mV in 10 mV steps for 250 ms) were applied and the whole-cell K^+^ current was recorded. The K^+^ channel blockers TEA (25 mM), clotrimazole (10 uM) and Ba^2+^ (5 mM) were applied to the cells using a perfusion pen.

## Results

Isolated duct cells express voltage-activated K^+^ channels. From a holding potential of −80 mV, K^+^ currents were activated at command voltages near 0 mV and reached amplitudes of 0.14 – 0.4 nA at +100 mV. Currents reached peak amplitude less than 20 ms after depolarization and did not inactivate. Response of representative cells are shown in Figures 3 and 4. TEA reversibly blocked a portion of this voltage-activated K^+^ current with 9 mM CaCl_2_ (1 µM free Ca^2+^) in the pipette solution (Fig. 3) in 6 out of 7 cells tested. The percentage of current block at +100 mV was (68 ± 17%, mean ± SD, n=6). This current was inactivated when the holding potential was kept at 0 mV in every cell tested (n=5, data not shown). Both Ca^2+^-dependent and Ca^2+^-independent K^+^ channels were present, and they had different responses to TEA. TEA had no effect on voltage-activated K^+^ currents in every cell tested (n=3) when Ca^2+^-free pipette solution was used (Fig. 4). Currents were not blocked by clotrimazole or Ba^2+^.

**Fig. 3.**
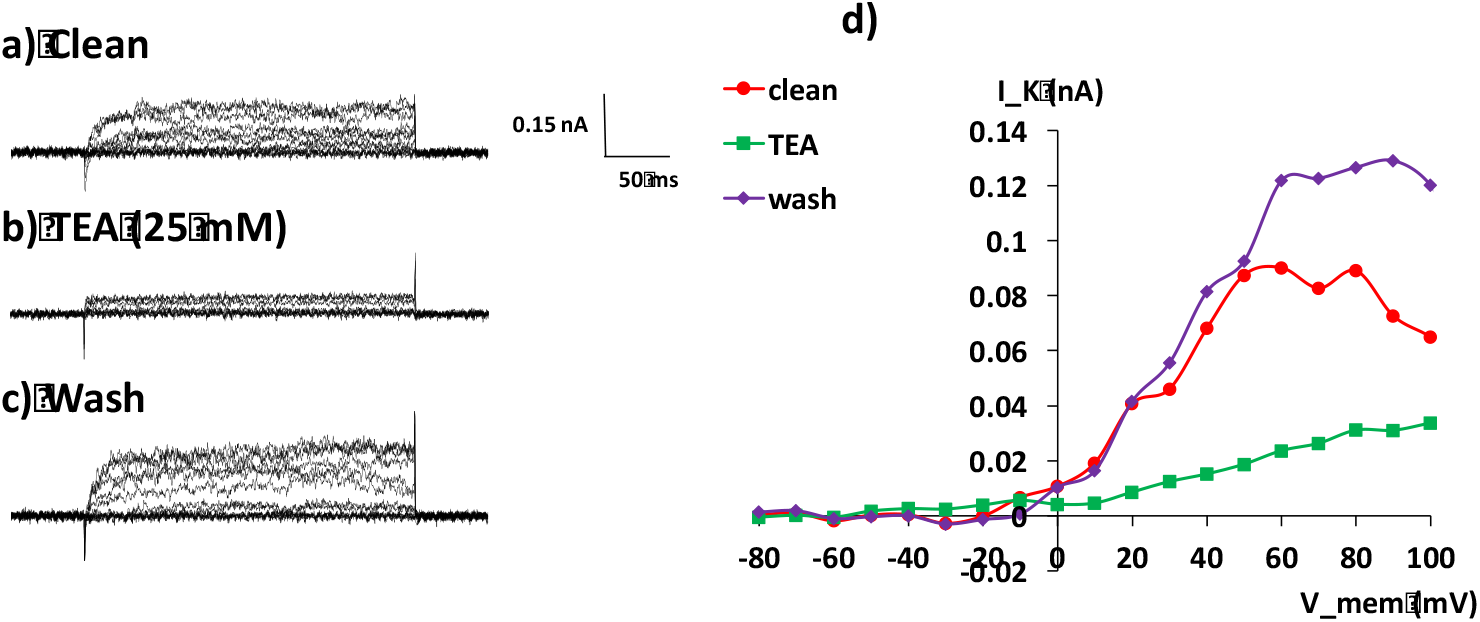
Voltage-activated K^+^ currents in duct cells are partially blocked by TEA when 9 mM CaCl_2_ (1 µM free Ca^2+^) is included in the pipette solution. From a holding potential of −80 mV, voltage steps of 250 ms duration were given in +10 mV steps, and whole-cell currents were recorded. a – c) Current traces from a representative duct cell with Ca^2+^ in pipette solution before TEA application, during TEA, and after washout. d) Current-voltage (I-V) relationships of the same cell.

**Fig. 4.**
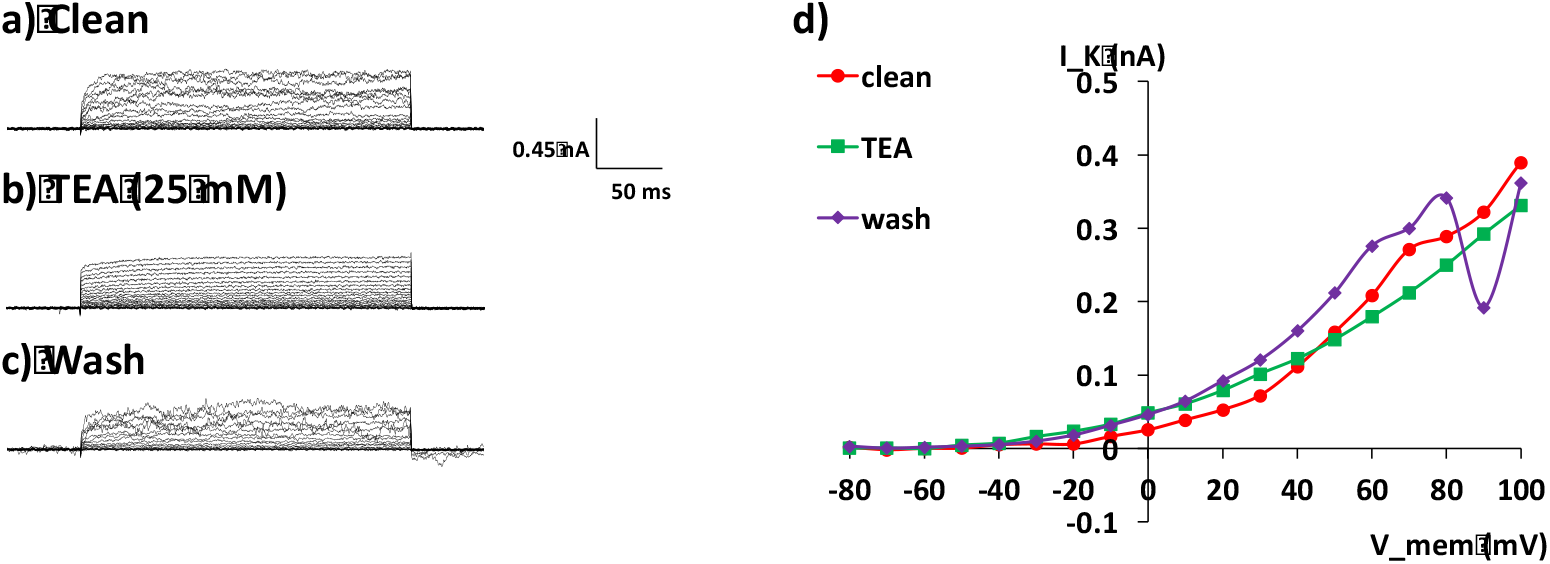
Voltage-activated K^+^ currents in duct cells are insensitive to TEA when Ca^2+^-free pipette solution is used. From a holding potential of −80 mV, voltage steps of 250 ms duration were given in +10 mV steps, and whole-cell currents were recorded. a – c) Current traces from a representative duct cell before TEA application, during TEA, and after washout. d) Current-voltage (I-V) relationships of the same cell.

As expected based on previous studies, isolated acinar cells also express voltage-activated K^+^ channels. TEA reversibly blocked nearly all voltage-activated K^+^ current with 9 mM CaCl_2_ (1 µM free Ca^2+^) in the pipette solution (Fig. 5) in every cell tested (n=12). In contrast to duct cells, however, TEA also blocked voltage-activated K^+^ currents in acinar cells in the absence of Ca^2+^ (data not shown).

**Fig. 5.**
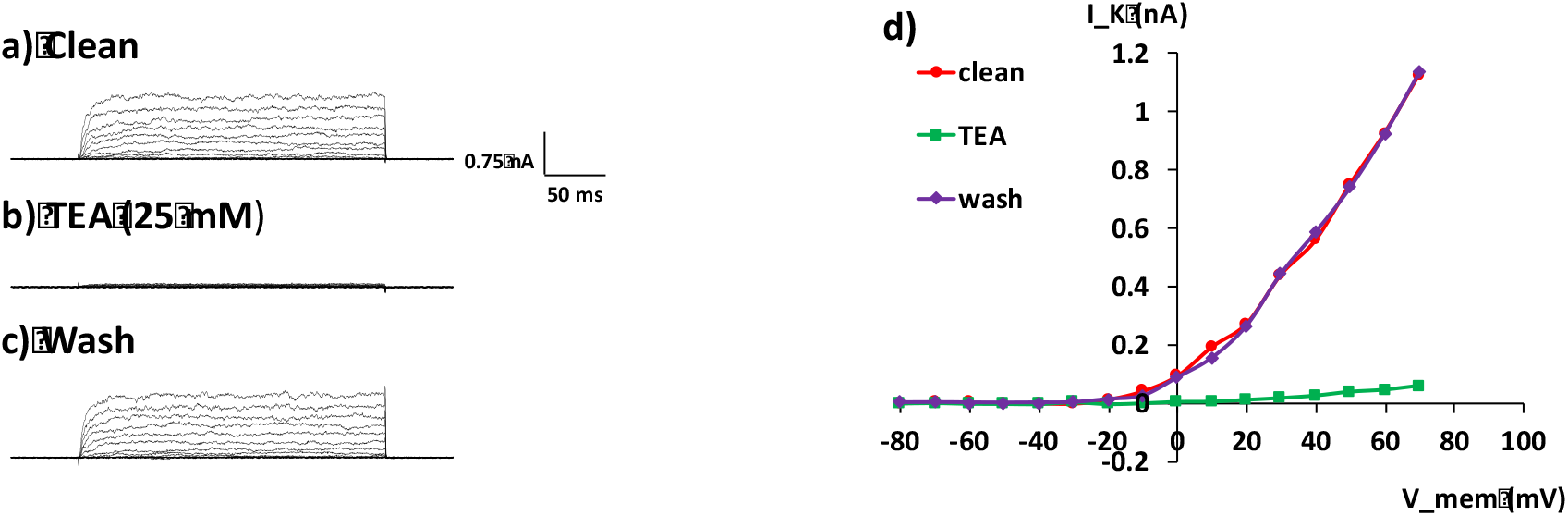
Voltage-activated K^+^ currents in acinar cells were reversibly blocked by TEA when 9 mM CaCl_2_ (1 µM free Ca^2+^) was included in the pipette solution. From a holding potential of −80 mV, voltage steps of 250 ms duration were given in +10 mV steps, and whole-cell currents were recorded. a – c) Current traces from a representative acinar cell before TEA application, during TEA, and after washout. d) Current-voltage (I-V) relationships of the same cell.

## Discussion

A method for isolating duct cells from rat lacrimal glands has been developed which provides viable duct cells for patch-clamp recording. While another brief report on K^+^ channels from cells in intact duct cells has been published,^14^ this is the first study giving a detailed description of the method for preparing single cells and is also the first compare ion channel currents in lacrimal gland duct cells to those in acinar cells. The method will be useful for those with an interest in further characterization of ion channel function in lacrimal gland duct cells.

Lacrimal gland duct cells have voltage-activated, Ca^2+^-dependent K^+^ channels that are sensitive to TEA, as well as voltage-activated Ca^2+^-independent K^+^ channels that are not blocked by TEA. In contrast to duct cells, TEA completely blocks voltage-activated K^+^ channels in acinar cells regardless of whether Ca^2+^ is included in the pipette.

The Ca^2+^-dependent K^+^ channel identified in this study probably corresponds to the IK_Ca_1 channel that we have identified on the apical membranes of rat lacrimal gland ducts.^7^ The presence of this channel, along with the apical KCC1 transporter and CFTR, which have been identified immunohistochemically and functionally,^7,9,10,15,16^ are consistent with evidence that the ducts secrete K^+^. The location of muscarinic receptors on the basal membrane of the duct cells,^7^ and the demonstration by Katona et al.^17^ that isolated ducts secrete fluid in response to carbachol, support a mechanism in which cholinergic stimulation causes release of intracellular calcium^18^ which activates IK_Ca_1 channels, resulting in secretion of fluid rich in K^+^. As we and others have reported,^19–21^ the high concentration of K^+^ in tears appears to be essential for the health of the ocular surface epithelium.

## Acknowledgements

Supported by NIH grant R01 EY 018100. Mark Schotanus, Susan Bardolph, Charlotte Stahl and Nhu Pham, provided technical assistance. The authors have no commercial relationships.

